# Cspg4 sculpts oligodendrocyte precursor cell morphology

**DOI:** 10.1101/2024.08.08.607226

**Authors:** Samantha Bromley-Coolidge, Diego Iruegas, Bruce Appel

## Abstract

The extracellular matrix (ECM) provides critical biochemical and structural cues that regulate neural development. Chondroitin sulfate proteoglycans (CSPGs), a major ECM component, have been implicated in modulating oligodendrocyte precursor cell (OPC) proliferation, migration, and maturation, but their specific roles in oligodendrocyte lineage cell (OLC) development and myelination *in vivo* remain poorly understood. Here, we use zebrafish as a model system to investigate the spatiotemporal dynamics of ECM deposition and CSPG localization during central nervous system (CNS) development, with a focus on their relationship to OLCs. We demonstrate that ECM components, including CSPGs, are dynamically expressed in distinct spatiotemporal patterns coinciding with OLC development and myelination. We found that zebrafish lacking *cspg4* function produced normal numbers of OLCs, which appeared to undergo proper differentiation. However, OPC morphology in mutant larvae was aberrant. Nevertheless, the number and length of myelin sheaths produced by mature oligodendrocytes were unaffected. These data indicate that Cspg4 regulates OPC morphogenesis *in vivo*, supporting the role of the ECM in neural development.

## Introduction

The extracellular matrix (ECM) is a complex network of proteins and proteoglycans that surrounds cells in all tissues, providing structural support and biochemical cues that regulate diverse cellular processes (Frantz et al., 2010). In the central nervous system (CNS), the neural ECM plays crucial roles in neural development (Barros et al., 2011; Melrose et al., 2021; Siebert & Osterhout, 2011), plasticity (Dzyubenko et al., 2021), and regeneration (reviewed in Sherman & Back, 2008).

Major components of the neural ECM are chondroitin sulfate proteoglycans (CSPGs), which consist of a core protein with covalently attached chondroitin sulfate glycosaminoglycan chains. CSPGs are highly expressed in the CNS and have been implicated in various neurodevelopmental processes, including neural stem cell proliferation (Sirko et al., 2007), axon guidance (Brittis et al., 2008; Kantor et al., 2004) synaptogenesis (Miyata et al., 2012; Pizzorusso et al., 2002), and myelination (Keough et al., 2016; Pendleton et al., 2013).

Myelination, the process by which oligodendrocytes ensheathe axons with a lipid-rich membrane, is essential for rapid propagation of action potentials (Huxley & Stampfli, 1948), metabolic support of axons (Fünfschilling et al., 2012; Nave, 2010) and proper neuronal function (Simons & Nave, 2016). Oligodendrocyte precursor cells (OPCs) are a distinct glial cell population that gives rise to myelinating oligodendrocytes (OLs) during development and throughout adulthood (Rivers et al., 2008; Zhu et al., 2011). The differentiation of OPCs into mature OLs is a tightly regulated process influenced by both intrinsic factors and extrinsic cues from the surrounding microenvironment (Emery, 2010).

Accumulating evidence suggests that the ECM plays a pivotal role in regulating OPC differentiation, migration, and myelination. During development, CSPGs have been shown to influence OPC migration and the timing of oligodendrocyte differentiation (Siebert & Osterhout, 2011; Szuchet et al., 2000). For instance, the CSPG Versican inhibits OPC differentiation and myelination in the optic nerve during early postnatal development (Dours-Zimmermann et al., 2009). In the context of CNS injury, CSPGs inhibit OPC differentiation and remyelination (Karus et al., 2016; Kuboyama et al., 2017). This could potentially occur through modulation of integrin and growth factor receptor signaling pathways, both of which interact with CSPGs (Chekenya et al., 2008; Fukushi et al., 2004; Lau et al., 2013; Siebert & Osterhout, 2011). Additionally, genetic ablation of CSPG receptors can enhance developmental myelination and remyelination (Pendleton et al., 2013; Shen et al., 2009). However, the specific mechanisms by which CSPGs influence OPC development, differentiation, and myelination in various contexts remain incompletely understood.

One CSPG that has attracted interest is the neural/glial antigen 2 (NG2) protein, encoded by the *Cspg4* gene. NG2 is a transmembrane CSPG that is highly expressed by OPCs in the developing and adult CNS (Akiko Nishiyama et al., 2009). *Cspg4* KO mice exhibit reduced OPC proliferation and delayed oligodendrocyte differentiation during development (K. Kucharova & Stallcup, 2010). These mice also show impaired remyelination following demyelinating injury, suggesting a critical role for NG2 in oligodendrocyte regeneration (Karolina Kucharova & Stallcup, 2015). Interestingly, *Cspg4* KO mice display altered synaptic plasticity and deficits in hippocampal long-term potentiation, indicating a broader role for NG2 in neuronal function (Sakry et al., 2014). *In vitro* studies have further elucidated the mechanisms by which NG2 influences OPC behavior. Many studies implicate NG2 in promoting cell proliferation and migration in response to growth factors such as PDGF and FGF2 (Górecki et al., 2007; Grako et al., 1999; Stallcup & Huang, 2008). Additionally, NG2-null OPCs exhibit impaired differentiation and remyelination *in vivo* (K. Kucharova & Stallcup, 2010; Karolina Kucharova et al., 2011). Mechanistically, NG2 has been shown to interact with various extracellular matrix components and growth factor receptors, modulating signaling pathways crucial for OPC proliferation, survival, and differentiation (Karolina Kucharova & Stallcup, 2015; Tamburini et al., 2019). Additionally, NG2 has been shown to promote mTOR-dependent translation, a critical mechanism for OPC growth and differentiation (Nayak et al., 2018).

Despite the prior studies of NG2/CSPG4 in OPC biology, its function during CNS myelination remains incompletely understood. To help fill this gap, we used zebrafish as a model system to investigate the specific contributions of NG2/CSPG4 to oligodendrocyte lineage cell development and myelination *in vivo*. We investigated the functional consequences of zebrafish *cspg4* mutations on OPC population dynamics, morphology, differentiation and subsequent myelination. Our findings provide insights into the interplay between the ECM, CSPGs, and oligodendrocyte lineage cells during CNS development and myelination, highlighting the importance of the NG2/CSPG4.

## Materials and Methods

### Animal husbandry

The Institutional Animal Care and Use Committee at the University of Colorado School of Medicine approved all animal work, in compliance with U.S. National Research Council’s Guide for the Care and Use of Laboratory Animals, the U.S. Public Health Service’s Policy on Humane Care and Use of Laboratory Animals, and Guide for the Care and Use of Laboratory Animals. Larvae were raised at 28.5°C in embryo medium (5 mM NaCl, 0.17 mM KCl, 0.33 mM CaCl, 0.33 mMMgSO4 (pH 7.4), with sodium bicarbonate) and staged as hours or days post fertilization (dpf) according to morphological criteria (Kimmel et al., 1995). Zebrafish lines used in this study included, AB wildtype, *cspg4^sa37999^* (ZDB-ALT-160601-5804), *cspg4^sa39470^* (ZDB-ALT-161003-16784), *Tg(mbpa:mCherry-CAAX)^co13Tg^* (A. N. Hughes & Appel, 2020). The *Tg(olig1:EGFP-CAAX)^co88Tg^* line was created by Natalie Carey using Gateway cloning to combine a 5.3 kb genomic fragment of zebrafish *olig1* into the construct *pEXPRS-olig1:EGFP-CAAX-pA2-Tol2,* which was injected into one cell stages embryos with mRNA encoding Tol2 transposase. EGFP+ embryos were raised to adulthood and outcrossed to identify founders that transmitted the transgene through the germline.

### Lectin fluorescence

2, 3, 5, and 7 dpf larvae were fixed in 4% paraformaldehyde in 1X PBS nutating overnight at 4°C. Post-fixation, larvae were rinsed in PBS, embedded in 1.5% agar/30% sucrose blocks, and then immersed in 30% sucrose overnight at 4°C. The blocks were subsequently frozen on dry ice and stored at –80°C. Transverse sections of 20 µm thickness were cut and collected on polarized microscope slides (Fisher Scientific 12-550-15). Sections were rehydrated with PBS and blocked for 1 hr at room temperature using a blocking solution (2% goat serum 2 mg/mL, 1uL of 1mg/mL unconjugated streptavidin (SNN1001 lot 2096048 in 1x PBS). The blocking solution is devoid of Triton X-100, which breaks down the ECM. Biotinylated *Vicia villosa* lectin (Vector Labs B1235-2 ZG0309) was then applied to the sections at 1:500 for 2 hrs at room temperature. Secondary antibody streptavidin-Cy5 (ThermoFisher SA1011) was applied 1:1000 O/N at 4°C. After washing with PBS, slides were mounted with Vectashield (Vector Laboratories H1000), coverslipped (VWR 48393-106), and dried overnight covered from light at 4°C. The next day, slides were sealed with clear nail polish and imaged in the subsequent 3 days.

### Bioinformatics

We used RNA-seq data from GEO dataset GSE52564, https://www.ncbi.nlm.nih.gov/geo/query/acc.cgi?acc=GSE52564 (Zhang et al., 2014).

We used the following genes to create a heatmap of RNA transcript levels: showing *Acan, Bcan, Cd44, Col10a1, Col11a1, Col11a2, Col12a1, Col13a1, Col14a1, Col15a1, Col16a1, Col17a1, Col18a1, Col19a1, Col1a1, Col1a2, Col20a1, Col22a1, Col23a1, Col24a1, Col25a1, Col26a1, Col27a1, Col28a1, Col2a1, Col3a1, Col4a1, Col4a2, Col4a3, Col4a3bp, Col4a4, Col4a5, Col4a6, Col5a1, Col5a2, Col5a3, Col6a1, Col6a2, Col6a3, Col6a4, Col6a5, Col6a6, Col7a1, Col8a1, Col8a2, Col9a1, Col9a2, Col9a3, Cspg4, Cspg5, Hspg2, Lama1, Lama2, Lama3, Lama4, Lama5, Lamb1, Lamb2, Lamb3, Lamc1, Lamc2, Lamc3, Ncan, Smc3, Tnc, Tnr, and Vcan*. Counts per million (CPM) values were calculated using the cpm function in the edgeR R package1 with log=F. Average values were calculated within their respective group: Astrocyte, microglia, myelinating oligodendrocytes, neuron, and oligodendrocyte precursor cells. Analysis was performed using Pluto (https://pluto.bio).

### Genotyping

For DNA extraction from zebrafish, two methods were utilized depending on the sample type. NaOH lysis was used for live-imaged zebrafish larvae aged 3-7 dpf, involving a brief incubation in a 50 mM NaOH solution for 10 min at 98°C to lyse cells and release DNA, vortexed for 10 sec, followed by neutralization with Tris-HCl. Proteinase K (Invitrogen AM2548) lysis was used for fixed tissues, following the manufacturer instructions. This method includes a heat inactivation step to preserve DNA integrity.

Genotyping of *cspg4^sa37999^* zebrafish samples used primers, [FWD:5’TCTGTGCCACTGATCATCGT3’, REV:5’GCAGCAGTGAATTTGAGGCT3’] to amplify a 290 bp fragment. The following thermocycler protocol was used for amplification: initial denaturation at 95°C for 3 minutes; followed by 35 cycles of 95°C for 30 seconds (denaturation), 54°C for 30 seconds (annealing), and 72°C for 1 minute (extension); then a final extension at 72°C for 5 minutes. Subsequently, 0.5uL XbaI (NEB R0145L) was added and incubated for 2 hours at 38°C, to selectively cleave the mutant fragment. The resulting fragments were then separated by electrophoresis on a 3% agarose gel to distinguish between mutant and wild-type alleles based on the presence or absence of one band at 290 bp indicating wildtype, one band around 150 bp (combination of 167 bp and 123 bp restriction digest products) indicating homozygous mutants, or two bands of 290 bp and 150 bp indicating heterozygotes.

Genotyping of *cspg4^sa39470^* zebrafish samples used primers, [FWD: 5’AAGGAGGGTCGACACTTACA3’, REV: 5’AATCTGACGGTGAGGGAAGG-3’] to amplify a 198 bp fragment. The following thermocycler protocol was used for amplification: initial denaturation at 95°C for 3 minutes; followed by 35 cycles of 95°C for 30 seconds (denaturation), 49°C for 30 seconds (annealing), and 72°C for 1 minute (extension); then a final extension at 72°C for 5 minutes. Subsequently, 0.5uL TSP45I (NEB R0583L) was added and incubated for 2 hours at 65°C, to selectively cleave the wild-type fragment. The resulting fragments were then separated by electrophoresis on a 3% agarose gel to distinguish between mutant and wild-type alleles based on the presence or absence of one band at 175 bp (23 bp runs off gel) indicating wildtype, one band at 198 bp indicating homozygous mutants, or two bands of 175 bp and 198 bp indicating heterozygotes.

PCR products were run next to a 100 bp ladder (Thermoscientific SM0243) on a 1.5% agarose A (BioPioneer C0009), 1.5% MetaPhor (Lonza 50184) ethidium bromide (Sigma-Aldrich 1239-45-8) electrophoresis gel. Images were obtained using a BioRad Gel Doc EZ Imager.

### Fluorescent *in situ* RNA hybridization

We performed *in situ* RNA hybridization using a RNAscope Multiplex Fluorescent v2 (ACD Bio 3231000) kit following the manufacturer’s protocol apart from the following: No Xylene, 15 uL slices, 30 min PFA post-fix rather than 2 hr. We used the following probes: Ch1 – *mbpa* (ACDbio 533861-C1), Ch2 – *sox10* (ACDbio 444691-C2), Ch3 – *cspg4* (ACDbio 529741-C3). Opal dyes used: 650 (FP1496001KT, lot 201008027:1, 520 (FP1487001KT, lot 210415025:1), 570 (FP1488001KT, lot 210408002:1). Slides were coverslipped with Vectashield (Vector Laboratories H1000), dried overnight at 4°C protected from light. The slides were then sealed with clear nail polish and imaged the next day. All slides were blinded for imaging and analysis.

### Mosaic labeling of OLCs

*Cspg4* mutant embryos were produced using crosses of heterozygote adults. Embryos were injected at the single cell stage with approximately 5 nL of a mixture containing 5 uL 0.4M KCl, 1 uL phenol red (Sigma-Aldrich P0290), 1 uL Tol2 RNA (250 ng/uL), 250 ng plasmid DNA in a final volume of 10 uL. We used plasmids *pEXPR.olig1:EGFP-CAAX-Tol2* to label OPCs and *pEXPR.mbpa:EGFP-CAAX-Tol2* to label OLs. We collected images of cells from living larvae at 3, 5, 7 dpf using confocal microscopy. Larvae were genotyped following image collection and analysis.

### Microscopy

We obtained microscope images of living larvae by embedding them laterally in 1.2% low-melt agarose (Agarose Unlimited PS1200) containing 0.4% tricaine (Western Chemicals NC0342409) for immobilization. We acquired images using a C-Apochromat 40x/1.1 NA water-immersion objectives on a Zeiss CellObserver Spinning Disk confocal system equipped with a Photometrics Prime 95B camera. For fluorescent *in situ* RNA hybridization images, we used a 40X/1.3 NA objective on the same microscope to image fixed tissue on slides.

### Image analysis

All images were blinded before analysis. For fluorescent *in situ* RNA hybridization only the middle 10 µm optical slices of each z-stack were analyzed using FIJI to determine how many cells expressed each RNA probe, then averaged per animal before comparing between conditions.

To perform OPC branching analysis, z-stack images were converted from czi to native Imaris format. OPC morphology was analyzed using the ‘Filament analysis’ tool with the following specifications: If there were any other cells or fluorescent signal in the z-stack, the cell for analysis was isolated via cropping the z-stack or using the masking tool, no loops method, largest diameter e.g. dendrite beginning 10 µm, smallest diameter e.g. dendrite ending 0.5 µm. Seed points were then established using the threshold tool and manually adding seed points to ensure the entire cell arbor was covered with approximately 1 µm between seed points, but importantly without overlapping seed points. Filaments were centered before gathering output data (including branch summed length, number of terminal points, Sholl analysis, and cellular territory using convex hull analysis). Data from multiple OPCs within one larva were averaged before comparing between groups.

To perform OL/myelin sheath morphology, z-stack images were converted from czi to native Imaris format. We used the ‘Filament analysis’ tool in the ‘skip automatic creation, edit manually’ setting. Filaments were added, one for each sheath, using ‘automatic depth’ viewing to trace each sheath individually in 3D. Output data included the number of sheaths, length of each sheath, and total summed length of sheaths per OL. Data from multiple OLs within one larva were averaged before comparing between groups.

### Statistics and data presentation

We performed statistical analyses using R, Version 2023.06.1+524 (2023.06.1+524) with the following libraries: asbio, dplyr, ggplot2, ggpubr, readxl, scales, tidyverse. We tested each dataset for normality using the Shapiro Test. Based on the output of the Shapiro Tests, the following statistical tests were used: OLC counts with RNAscope – ANOVA and Tukey post hoc; OPC summed process length– Kruskal-Wallis; OPC Sholl analysis – Kruskal-Wallis chi square test and Dunn test with Bonferroni correction; OL/myelin morphology – Kruskal-Wallis.

Gene expression heatmaps were created with Pluto (https://pluto.bio). All other plots were created in R, Version 2023.06.1+524 (2023.06.1+524) using the following libraries: asbio, dplyr, ggplot2, ggpubr, readxl, scales, tidyverse.

## Results

### The spinal cord ECM develops in a spatially and temporally specific pattern

As a first step toward understanding how the ECM might guide oligodendrocyte lineage cell (OLC) development and myelination, we employed the use of lectin *vicia villosa lectin (*VVL). This lectin, which is commonly used to visualize neural ECM (Derouiche et al., 1996; Ojima et al., 1998), specifically labels N-acetylgalactosamine residues. These residues are enriched on chondroitin sulfate proteoglycans (CSPGs), which are important functional contributors to the ECM and highly expressed by OLCs (Asher et al., 2002; A. Nishiyama et al., 1996). To label OLCs at different stages of maturity in combination with ECM, we stained tissue sections from larvae carrying the transgenes *Tg(olig1:EGFP-CAAX)* and *Tg(mbpa:mCherry-CAAX),* which mark OPCs and OLs respectively, with VVL.

Representative images from 3-7 days post fertilization (dpf) of transverse spinal cord sections revealed pervasive VVL labeling throughout the developing CNS, resembling the honeycomb-like structures described in other studies (Horii-Hayashi et al., 2015; Murakami, 1994; Paul & Ulfig, 1998) (Figure 1A-C). Throughout the time course, distinct puncta were evident in both developing axon- and soma-rich regions. Notably, we observed a high deposition of VVL staining at the dorsal edge of the dorsal spinal white matter (WM) tract in 5 dpf larvae (Figure 1B). This deposition was adjacent to the most dorsal OLCs. The specific enrichment of VVL staining in this region suggests a potential role for lectin-labeled CSPGs in the interaction between OLCs and the developing dorsal spinal WM tract. By 7 dpf, VVL staining showed further refinement, with pronounced enrichment in the axon-rich regions (Figure 1C). These temporal changes in VVL staining patterns informed our choice of timepoints for subsequent morphological analyses, allowing us to capture key stages in the interaction between CSPGs and OLC development.

**Figure 1.**
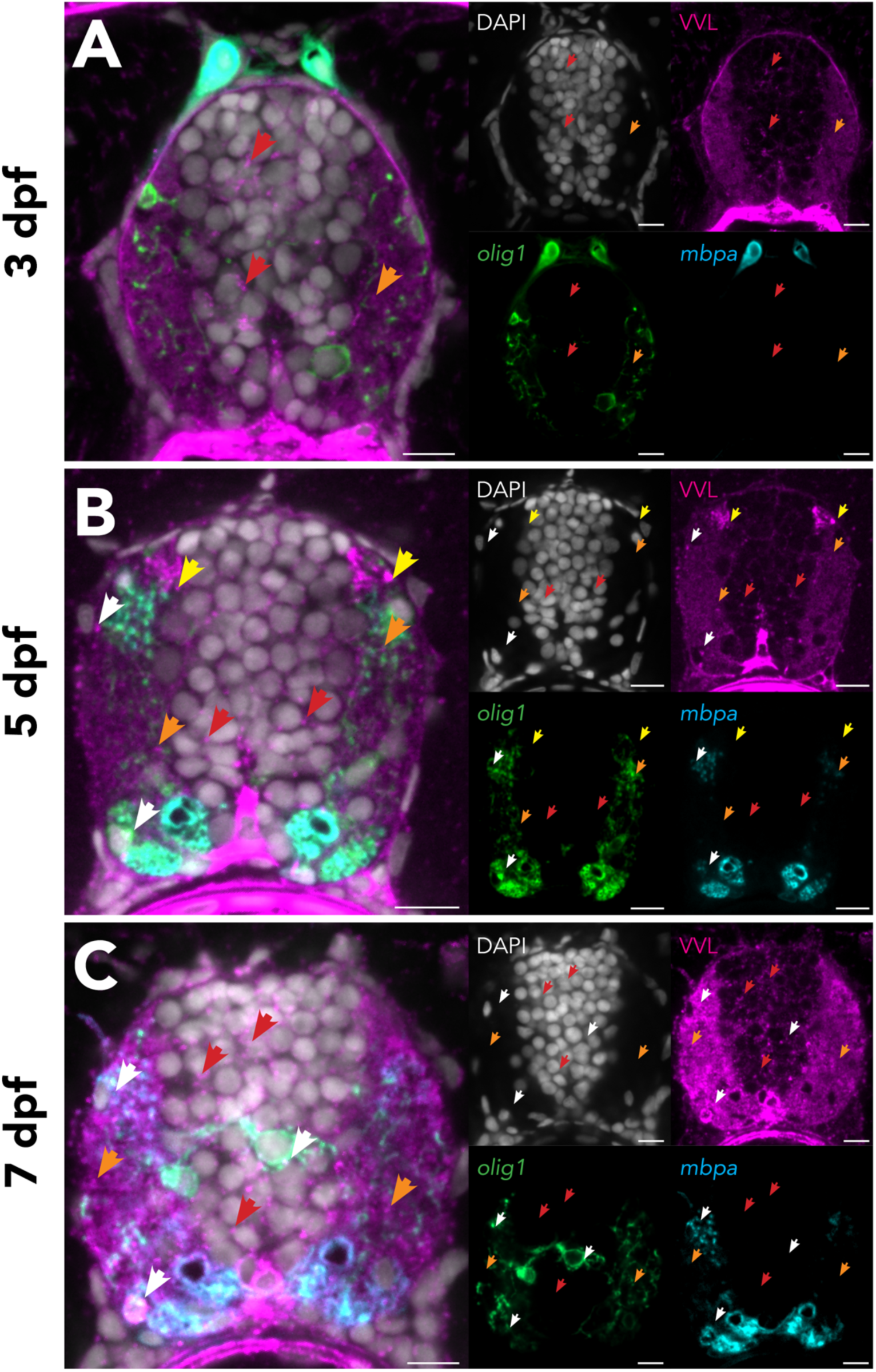
Lectin staining intensity increases in the spinal cord during zebrafish development and is dynamically located. Transverse slices of zebrafish larval spinal cord of larvae containing transgenes *Tg(olig1:EGFP-CAAX)* and *Tg(mbpa:mCherry-CAAX)* at 3, 5, and 7 days dpf. Arrowheads indicate specific fluorescence patterns of *vicia villosa* lectin: red arrowheads highlight puncta in soma-rich areas; orange arrowheads highlight puncta in axon-rich areas; white arrowheads highlight puncta near oligodendrocyte lineage cells (OLCs); and yellow arrowheads highlight strong deposition at the dorsal white matter tract. Scale bars 10 µm.

### OPCs express many ECM genes, including *cspg4*

To investigate the expression of genes encoding ECM components across different cell types in the CNS, we analyzed transcriptomic data from mouse cerebral cortex (Zhang et al., 2014). We focused on structural ECM genes, including chondroitin sulfate proteoglycans (*CSPGs*), heparan sulfate proteoglycans (*HSPGs*), collagens, laminins, and tenascins.

Our analysis revealed that OPCs express numerous ECM-encoding genes at high levels (Figure 2A, B). Notably, myelinating OLs express much lower levels of ECM genes, suggesting that transcription of ECM-encoding genes is downregulated concomitant with OL differentiation. These data indicate that, together with astrocytes, OPCs produce a large amount of the CNS ECM.

**Figure 2.**
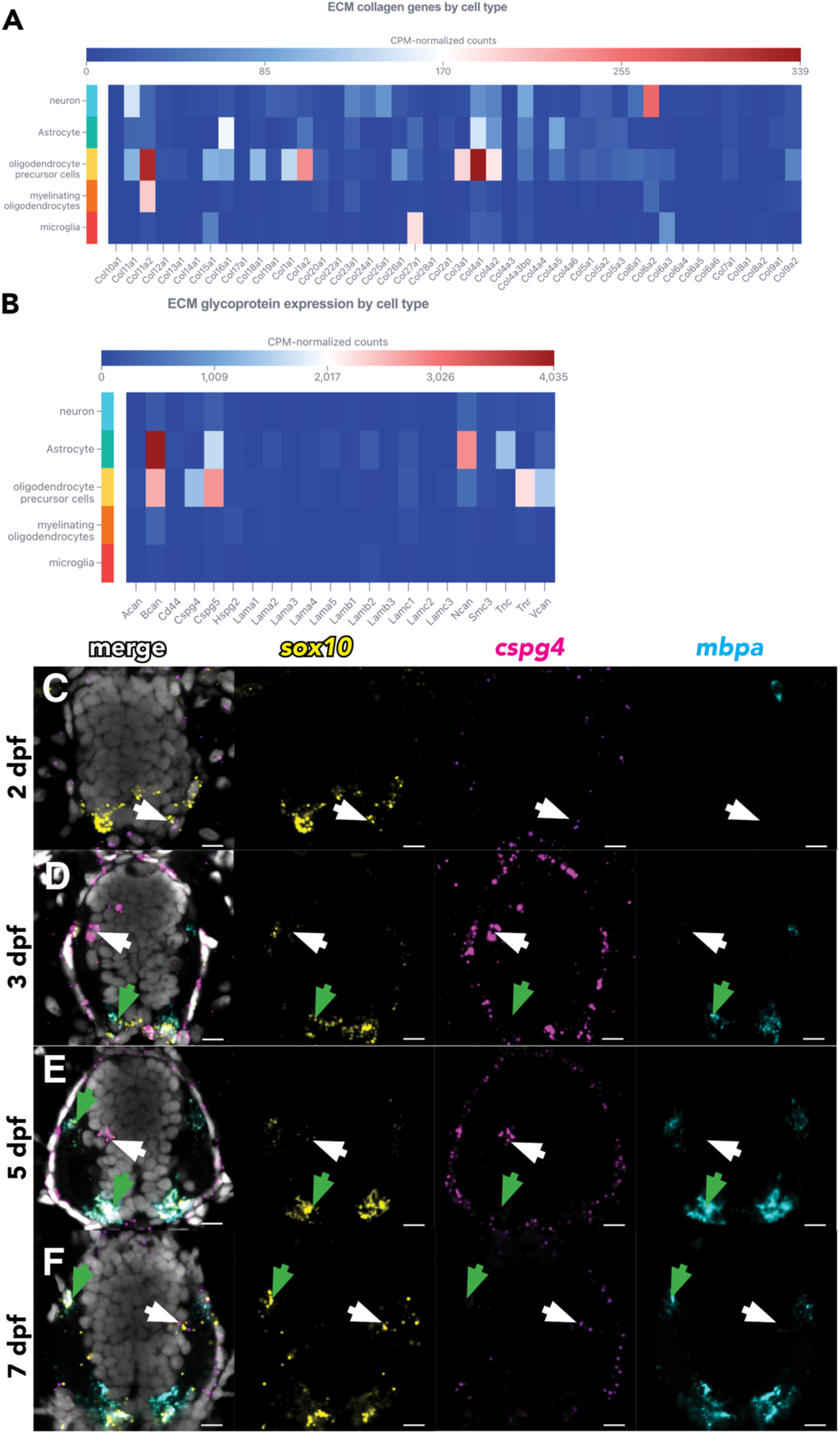
OPCs and astrocytes have highest expression of structural ECM genes, and zebrafish OPCs express *cspg4*. Heatmaps showing expression levels of structural extracellular matrix (ECM) genes – collagens (A), glycoproteins (B) – across cell populations in mouse cortex. Dataset from Zhang et al, 2014. Heatmaps generated using pluto.com. Transverse slices of wild-type zebrafish larval spinal cord at 2 (C), 3 (D), 5 (E), and 7 (F) dpf processed to detect *sox10* (yellow), *cspg4* (magenta), and *mbpa* (cyan) transcripts. Nuclei are labeled with DAPI (grey). White arrows highlight OPCs co-expressing *sox10* and *cspg4*. Green arrows indicate OLs expressing *sox10* and *mbpa* but not *cspg4*. The left column shows merged imaged of all 4 channels. The other columns show individual images of *sox10*, *cspg4,* and *mbpa* expression. Scale bars 10 µm.

These data also indicated that, whereas many ECM genes expressed by OPCs are also expressed by astrocytes and neurons, *Cspg4* expression appears to be exclusive to OPCs. To validate this conclusion in zebrafish, we used fluorescent *in situ* RNA hybridization to detect *cspg4* transcripts in combination with *sox10* RNA, which marks all OLCs, and *mbpa*, which marks OLs. In zebrafish, spinal cord *sox10* expression is first evident at approximately 36 hours postfertilization (hpf), marking initiation of OPC formation. At 2 dpf, *sox10*+ OPCs express little *cspg4* (Figure 2D). At 3, 5, and 7 dpf, *cspg4* is evident in *sox10+ mbpa–* OPCs but absent from *sox10+ mbpa+* OLs (Figure 2D-F). These data are consistent with the RNA-seq data indicating that OPCs express *cspg4* and downregulate *cspg4* transcription as they differentiate as OLs.

### Loss of *cspg4* function does not impair OLC number or differentiation

To investigate *cspg4* function, we procured two mutant lines, *cspg4^sa37999^* and *cspg4^sa39470^*, with distinct point mutations affecting the *cspg4* gene (Figure 3A). The *cspg4^sa37999^* allele is an A to T mutation that introduces a premature stop codon early in exon 3. The *cspg4^sa39470^* allele is a C to T mutation at the splice donor site of exon 4, which is predicted to cause intron retention and introduction of a premature stop codon. We confirmed these alleles using PCR-based genotyping (Figure 3B).

**Figure 3.**
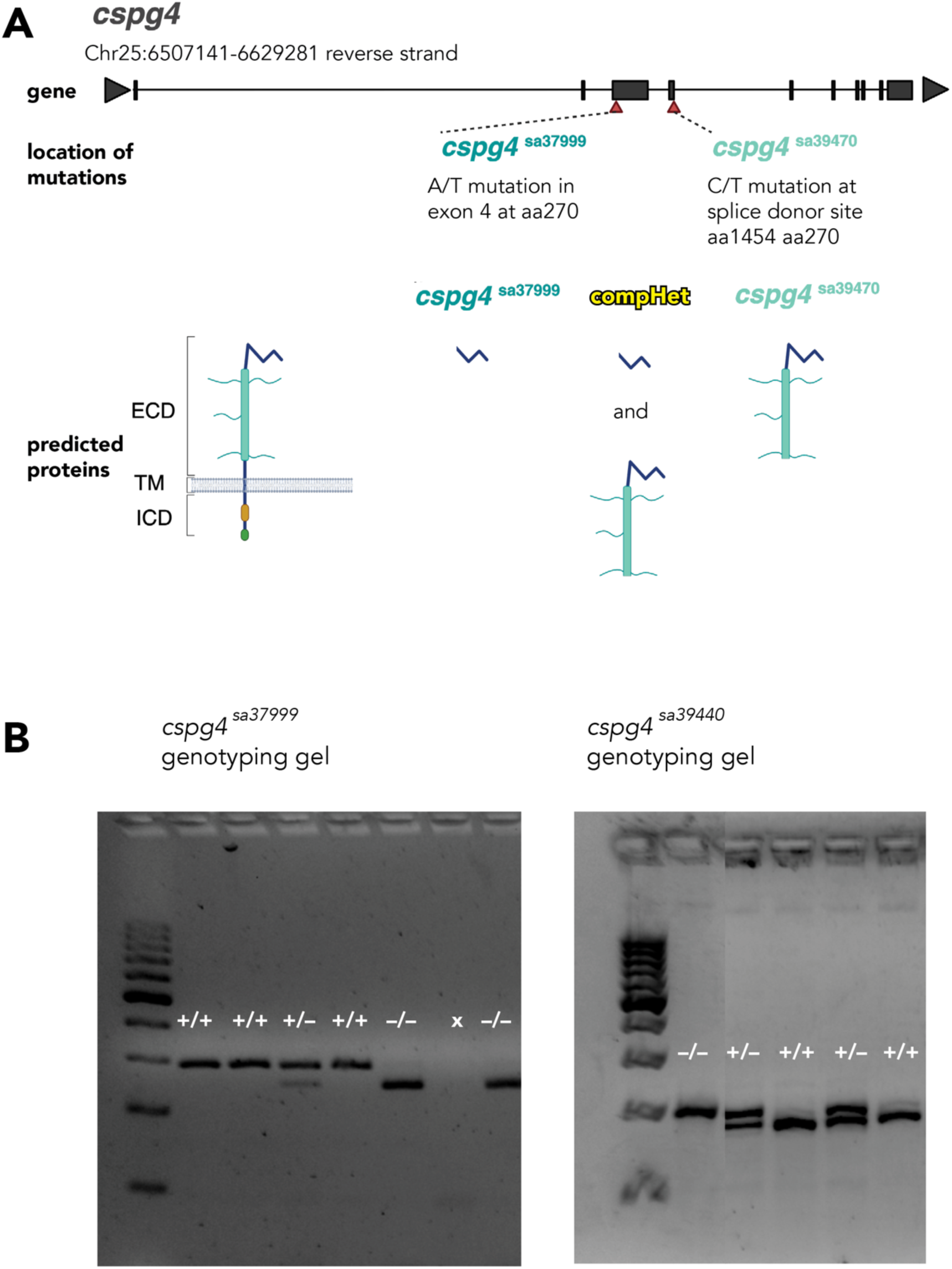
Schematic overview and confirmation of *cspg4* mutant alleles. (A) Locations of mutant alleles indicated by red arrowheads. Predicted proteins indicated below. compHet represents compound heterozygous *cspg4^sa37999^/*cspg4sa39470. Abbreviations: extracellular domain (ECD), transmembrane (TM), intracellular domain (ICD). (B) Genotyping results for PCR-amplified regions displayed on an agarose gel. For *cspg4^sa37999^*: 290 bp band indicates wildtype, ∼150 bp band (split into 167 bp and 123 bp) indicates homozygous mutation (−/−), and dual bands indicate heterozygous (+/−). For *cspg4^sa39470^*: 175 bp band signifies wildtype, 198 bp indicates homozygous mutation (−/−), and presence of both bands indicates heterozygosity (+/−).

To determine if loss of *cspg4* function affects OLC formation, we utilized fluorescent *in situ* RNA hybridization to visualize *sox10* and *mbpa* in transverse sections of wild-type and *cspg4^sa37999^* homozygous mutant embryos and larvae. We compared the numbers of *sox10+ mbpa–* OPCs and *sox10+ mbpa+* OLs from 3 to 7 dpf. Neither OPCs nor OLs differed significantly in number between genotypes although 2 dpf *cspg4^sa37999^* homozygous mutant embryos trended toward more OPCs than wildtype (p = 0.05). We conclude that loss of *cspg4* function does not interfere with OPC specification and OL differentiation.

**Figure 4.**
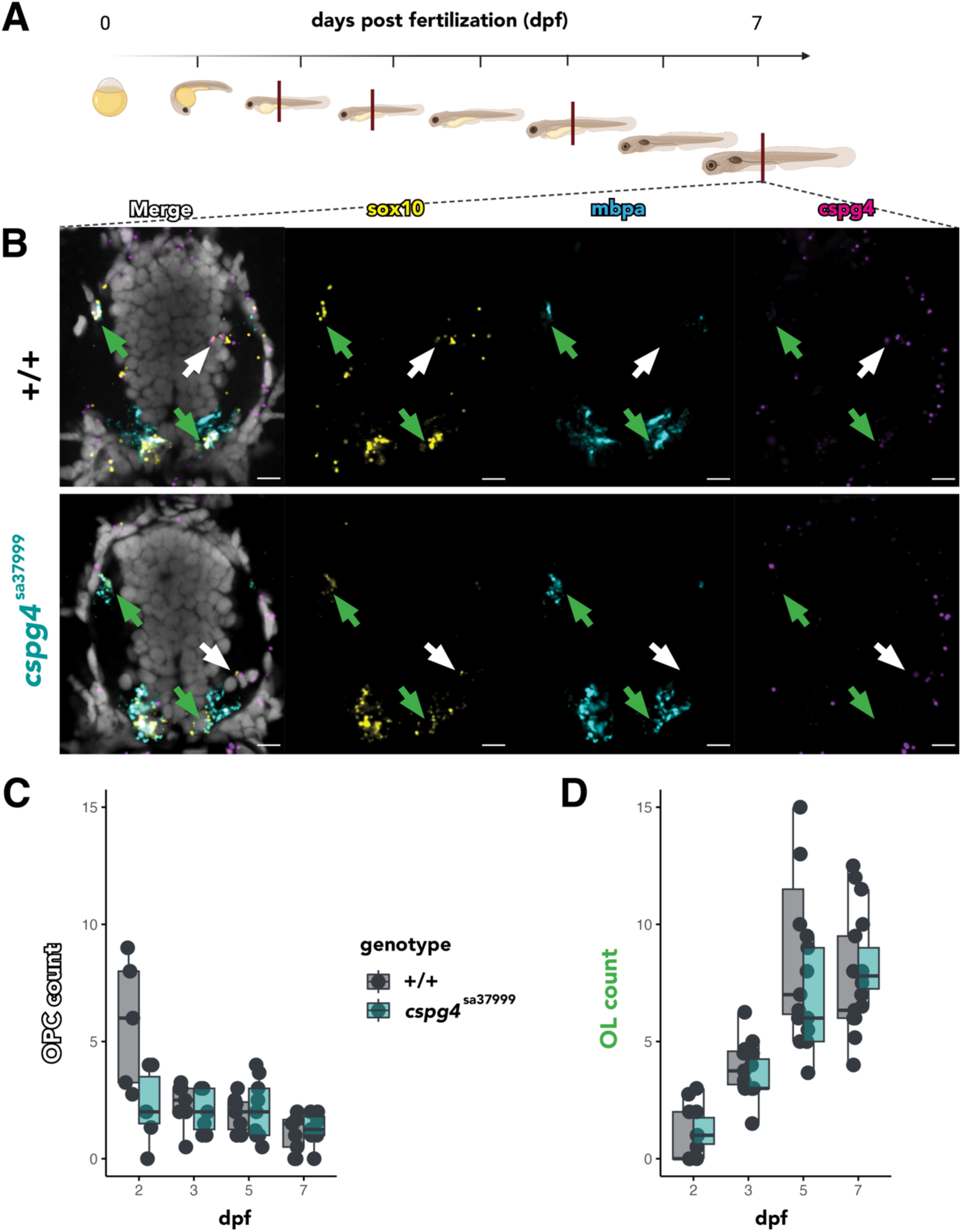
Loss of *cspg4* function does not impair OPC formation or OL differentiation. (A) Schematic representation of the experiment wherein larvae were fixed and sectioned at 2, 3, 5, and 7 dpf. Transverse sections were only analyzed if they showed the spine over yolk tube, indicated by crimson vertical lines. (B) Representative images of wild-type and *cspg4^sa37999^* homozygous mutant larvae showing *sox10* (yellow), *mbpa* (blue), and *cspg4* (magenta) RNAs detected using fluorescent *in situ* RNA hybridization; all cell nuclei labeled using DAPI (grey). White arrowheads highlight *sox10*+ *mbpa–* OPCs; green arrowheads highlight *sox10+ mbpa+* OLs. Scale bars all 10 µm. (C) Box plot of the number of *sox10+ mbpa-* OPCs per larva at each timepoint. (D) Box plot of the number of *sox10+ mbpa+* OLs at each timepoint. OPC and OL numbers were not statistically different between genotypes, although mutant embryos trended toward fewer OPCs at 2 dpf (p=0.054). Data analyzed by Kruskal-Wallis statistical test.

### OPCs are more complex in *cspg4* mutant zebrafish larvae

Previous studies have shown that CSPG4 interacts with ligands and structural extracellular matrix proteins (Burg et al., 1997; Goretzki et al., 1999; Tillet et al., 1997), raising the possibility that it conveys morphological cues or otherwise affects cellular complexity. To investigate whether loss of *cspg4* function alters OPC morphology in vivo, we injected single cell stage embryos with *olig1:EGFP-CAAX* plasmid, which marks OPCs with membrane-tethered EGFP, with synthetic mRNA encoding Tol2 transposase. We assessed OPCs in the living larvae using confocal microscopy. We focused our investigation on the spinal region positioned above the yolk tube to limit the effect of position along the rostro-caudal axis on maturation.

Representative images of fluorescently labeled OPCs are depicted in Figure 5A. We implemented a 3D Sholl analysis to give a detailed readout of complexity of the OPC branching arbors. This method uses concentric spheres originating at the soma and calculates the number of intersections with branches at 1 um intervals from the soma. Our results reveal that from 3 to 5 dpf, cells across all genotypes increased the total length of processes produced by individual OPCs, suggesting that they grew in size and complexity (Figure 5B). Interestingly, between 5 to 7 dpf, total process length regressed to 3 dpf levels in all conditions (Figure 5B).

**Figure 5.**
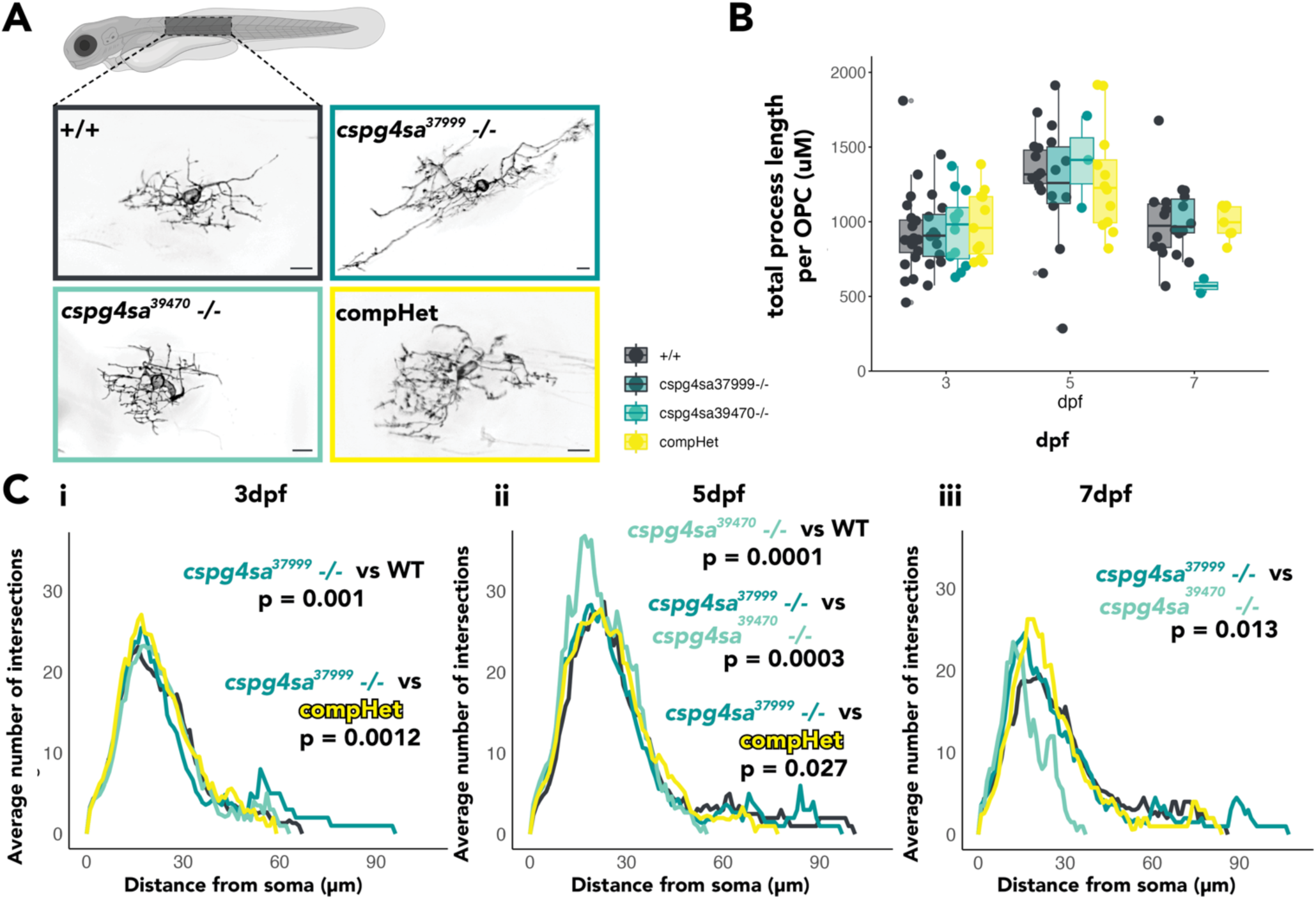
OPCs from *cspg4* mutant larvae have morphological differences in complexity compared to WT through development. (A) Schema of the imaging region in the spine above the yolk tube in larvae with example images of mosaically labeled fluorescent OPCs using *olig1:EGFPcaax*, imaged in live larvae. Color coding for genotypes is consistent across all figures. Scale bars all 10 µm. (B) The summed filament length of mutant larvae are not significantly different than WT at any timepoint using a two-way, non-repeated measures ANOVA. (C) Morphological complexity was assessed by Sholl analysis and Kruskal-Wallis test followed by a Dunn test when significant. This showed *cspg4^sa37999^* mutants have at least one significantly longer process on average than comHets (p = 0.001) at 3 dpf (i). *cspg4^sa39470^* mutants have significantly smaller cells with higher complexity closer to the soma compared to WT (p = 0.0001) at 5dpf (ii). At 7dpf (iii), *cspg4^sa37999^* mutants may be more complex close to the soma or have at least one longer process compared to WT (p=0.056).

We compared the number of intersections made by processes of individual OPCs across different genotypes at 3, 5, and 7 dpf. At 3 dpf, OPCs in homozygous *cspg4^sa37999^* mutant larvae showed significantly longer processes compared to wildtype (p = 0.001) and compound heterozygous (compHet) (p = 0.0012) larvae, while *cspg4^sa39470^* and compHet OPCs were not significantly different from wildtype (Figure 5Ci). By 5 dpf, a distinct phenotype emerged in *cspg4^sa39470^* homozygous mutants, with OPCs displaying significantly higher complexity near the soma but shorter processes overall compared to all other groups (wildtype p = 0.0001, *cspg4^sa37999^* p = 0.0003, compHet p = 0.027) (Figure 5Cii). At 7 dpf, all mutant alleles exhibited higher peaks in their Sholl traces compared to wildtype, indicating increased arbor complexity close to the cell soma. Notably, *cspg4^sa37999^* and *cspg4^sa39470^* homozygous mutants showed significantly different morphologies (p = 0.013), with *cspg4^sa37999^* OPCs maintaining longer processes and *cspg4^sa39470^* OPCs displaying shorter processes (Figure 5Ciii). These snapshots of OPC arbor morphology across this critical period of developmental OLC expansion and myelination reveal that each homozygous mutant allele yields distinct alterations in OPC morphology. While *cspg4^sa37999^* OPCs initially show longer processes, by 7 dpf, all mutant alleles develop a “bushy” rather than “tree-like” arbor complexity close to the soma. Furthermore, the two homozygous alleles exhibit opposing phenotypes in terms of process length, with *cspg4^sa37999^* OPCs maintaining longer processes and *cspg4^sa39470^* OPCs showing truncated processes. These morphological changes in *cspg4* mutant OPCs suggest altered cellular architecture, which could potentially impact their interactions with the surrounding extracellular matrix and neighboring cells.

### OLs in mutant larvae make myelin sheaths of similar number and length

To investigate whether loss of *cspg4* function alters myelin sheath formation, we injected single cell stage embryos with the plasmid *mbpa:EGFP-CAAX*, which labels OLs with membrane-tethered EGFP, and used confocal microscopy to assess the number and cumulative length of myelin sheaths formed by individual cells. This revealed that OLs in mutant larvae had morphologies comparable to those of wild-type controls (Figure 6A). Specifically, there were no significant differences in the number of myelin sheaths per OL (Figure 6B) or the total summed length of these sheaths between mutant and wild-type larvae (Figure 6C). Thus, loss of *cspg4* function does not appear to interfere with the myelinating capacity of oligodendrocytes.

**Figure 6.**
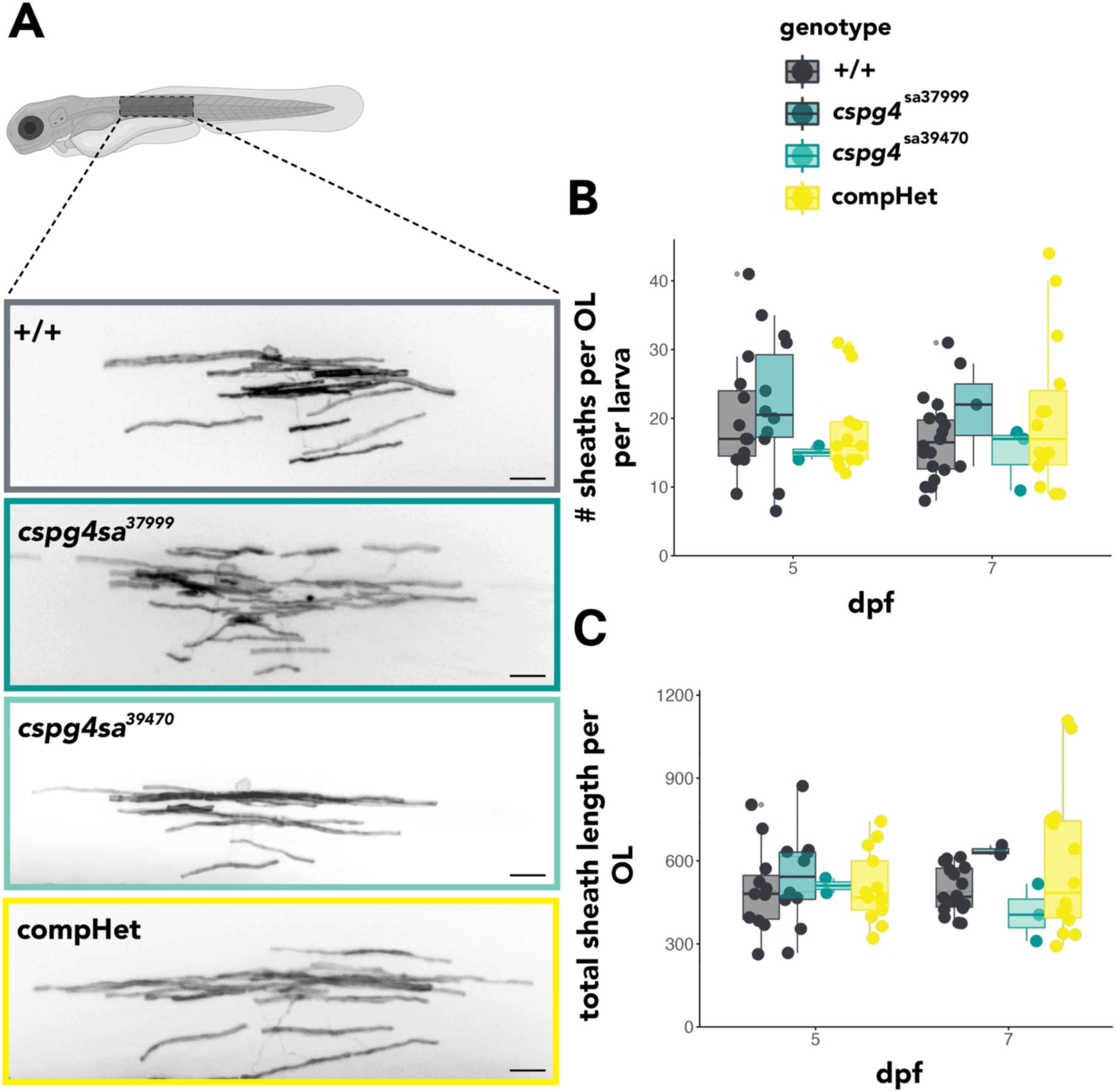
Loss of *cspg4* does not alter myelin sheath number or length. (A) Schema of the imaging region in the spine above the yolk tube in larvae with example images of mosaically labeled fluorescent OLs using *mbpa:EGFP-CAAX*. Using a Kruskal-Wallis statistical test, mutant larvae have oligodendrocytes with similar sheath number (B) and total summed length (C) compared to wildtype. Scale bars equal 10 µm.

## Discussion

Numerous genes, expressed by both neurons and glia, contribute to the extensive ECM of the CNS. In this context, CSPG4/NG2 is remarkably specific to OPCs. This specificity has been useful for investigation of OPCs in rodents, using immunohistochemistry to detect NG2 and *Cspg4* transgenes to identify and manipulate OPCs *in vivo*. However, there is much less information about zebrafish CNS *cspg4* expression. Here, we carefully analyzed *cspg4* expression through early larval stages using a highly sensitive fluorescent *in situ* RNA hybridization technique. This showed that zebrafish OPCs but not OLs or other spinal cord cells express *cspg4*. Thus, OPC expression of *cspg4* is conserved between rodents and zebrafish.

The first main finding from our efforts to understand CSPG4 function is that the number of OPCs in *cspg4*-deficient zebrafish larvae is not substantially different from wildtype. This is in contrast to *Cspg4* mutant mice, which have fewer OPCs in developing cerebellar white matter than wild-type mice; however, it is in agreement with other regions of the *Cspg4* knockout mouse, in which OPC number and development were not significantly affected (Hill et al., 2011; K. Kucharova & Stallcup, 2010). CSPG4 interacts with PDGFRa on OLCs (A. Nishiyama et al., 1996) and PDGF is a potent OLC mitogen *in vitro* (Richardson et al., 1988) and *in vivo* (Fruttiger et al., 1999). Therefore, in mice, CSPG4 might accentuate growth factor driven proliferation. What might explain the absence of a proliferative effect in zebrafish *cspg4* mutant larvae? Notably, larval zebrafish OPCs express *pdgfra* at very low levels (Sur et al., 2023) and zebrafish OPCs divide infrequently in early larval stages (Snyder et al., 2012). This raises the possibility that there is a relatively low level of growth factor driven OPC proliferation at early developmental stages that may underlie the difference in observations.

Our next main finding was that zebrafish *cspg4* mutant larvae have abnormal OPC morphology. *In vivo*, OPC morphologies are remarkably complex and dynamic (E. G. Hughes et al., 2013; Kirby et al., 2006; Marisca et al., 2020). Conceivably, interactions with the ECM contributes to these characteristics. By measuring OPC process length, number, and number of branches – binned by distance from the soma – as indicators of cellular complexity, we found that OPCs in *cspg4* mutant larvae generally had more complex morphologies than OPCs in wild-type larvae, resulting from changes in process length and the number of branches at various distances from the soma. Notably, we also found differences between the two mutant alleles and the compound heterozygotes, potentially reflecting different effects of the truncated proteins, which differ in the lengths of their extracellular domains. The *cspg4^sa37999^* allele, which is predicted to produce only the LGR binding domains, lacks critical components such as the chondroitin sulfate GAG (glycosoaminoglycan) side-chain domain, proteolytic cleavage sites, transmembrane domains, and the intracellular domain, which contains a PDZ interacting motif (Rolih et al., 2017). The *cspg4^sa39470^* allele contains the domains of the *cspg4^sa37999^* allele, with the addition of GAG side-chains, FGF and PDGFR binding domains. The presence or absence of each domain, and resulting cellular morphologies, may point to key signaling pathways that underly the morphological phenotypes. Future studies dissecting the specific contributions of each domain to OLC development will be crucial for understanding the multifaceted roles of CSPG4 *in vivo*.

Despite differences in OPC morphology, the number and length of myelin sheaths formed by OLs in *cspg4* mutant larvae were similar to those in wild-type larvae. Additionally, *cspg4* mutant larvae expressed *mbpa*, a marker of OL differentiation and myelination, similarly to wildtype. Thus, OPCs lacking *cspg4* are able to undergo normal differentiation as myelinating OLs. Although myelination was delayed in *Cspg4* mutant mice (K. Kucharova & Stallcup, 2010), the delay most likely reflected a deficit in initial OPC production, which we did not find in zebrafish.

OPCs are key players in CNS development, homeostasis, and repair (reviewed in Bergles & Richardson, 2016). OPCs provide trophic support to neurons and axons through the secretion of growth factors such as brain-derived neurotrophic factor (BDNF) and insulin-like growth factor 1 (IGF-1) (Wilkins et al., 2001, 2003). OPCs also modulate synaptic circuitry and plasticity through direct contacts with neurons and the release of signaling molecules (Sakry et al., 2014; Xiao et al., 2022), and contribute to blood-brain barrier integrity and function through interactions with endothelial cells (Seo et al., 2014). Importantly, missense mutations of human *CSPG4* have been linked to schizophrenia (de Vrij et al., 2019). Potentially, the changes in OPC morphology resulting from loss of CSPG4 function that we report here could impact how OPCs contribute to neural function.

It is important to acknowledge limitations of our study. First, CSPG4 is only one of numerous ECM components of the nervous system. Many of these molecules might contribute similarly to tissue microenvironments and intercellular signal transduction. For example, OPCs also express large amounts of CSPG5 (Zhang et al., 2014), which is closely similar to CSPG4 but has not been extensively studied. Therefore, simultaneous elimination of CSPG4 and CSPG5 might result in much stronger effects on OPC morphology. Second, our investigation focused only on how loss of CSPG4 function affects features of OPCs and OLs. Future studies will be required to assess the roles of CSGP4 produced by OPCs in neural circuit formation and function.

In conclusion, our study provides an *in vivo* analysis of *cspg4* function in sculpting OLC development. Our findings contribute to understanding the spatiotemporal dynamics of ECM-OLC interactions and the consequences of *cspg4* perturbation, which help advance our basic understanding of neurodevelopmental processes.

## Acknowledgments.

We thank Natalie Carey for creating and sharing the *Tg(olig1:EGFP-CAAX)* line and Dr. Julie Siegenthaler for comments on the manuscript.

## Declaration of interests

none

## Author contributions

Samantha Bromley-Coolidge: conceptualization, formal analysis, investigation, visualization, writing – original draft.

Diego Iruegas: investigation, formal analysis.

Bruce Appel: conceptualization, funding acquisition, project administration, writing – review and editing.

## Funding

This work was supported by the National Institutes of Health (NS406668 to B.A.) and a gift from the Gates Frontiers Fund to B.A.

## Abbreviations

OPC: oligodendrocyte precursor cell
OLC: oligodendrocyte lineage cell
OL: oligodendrocyte
CNS: central nervous system
ECM: extracellular matrix
CSPG: chondroitin sulfate proteoglycan
hpf: hours postfertilization
dpf: days postfertilization
GAG: glycosoaminoglycan

## Notes

### Competing Interest Statement

The authors have declared no competing interest.

